# Quantitative Magnetic Flow Cytometry in High Hematocrit Conditions for Point-of-Care Testing

**DOI:** 10.1101/2024.06.11.598398

**Authors:** Moritz Leuthner, Michael Helou, Mathias Reisbeck, Oliver Hayden

## Abstract

Quantitative cell analysis in liquid biopsies is essential for many clinical decisions, but it is primarily tied to centralized laboratories. However, access to these laboratories is limited in low-resource settings or for immobile patients, highlighting the urgent need for Point-of-Care (POC) testing infrastructure. Magnetic flow cytometers (MFC) offer a solution, albeit sample processing steps like cell lysis or washing crucially disrupt POC-capable MFC workflows. Here, we investigate conditions for immunomagnetic labeling and direct cell quantification in a streamlined workflow suitable for high hematocrit environments. Magnetic nanoparticles (MNP) are characterized by their size, magnetic moment, and potential to generate signal noise, favoring small (< 50 nm) MNPs. Theoretical models provide the framework for quantifying bound MNPs per cell, revealing labeling quality and giving insight into system requirements for reliable cell detection. Temporal labeling dynamics show suboptimal binding kinetics in whole blood (WB), leading to long incubation periods and only 50% recovery of optically determined concentrations. Besides showing quantitative MFC in WB with biomimetic microbeads, we finally quantify CD14^+^ monocytes in WB with our streamlined workflow, achieving an intra-assay coefficient of variation (CV) of 0.11 and a CV across multiple donors of 0.10, demonstrating reliable POC flow cytometry close to regulatory standards.

## 1. Introduction

The Point-of-Care (POC) testing landscape has experienced an enormous rise in awareness in the last few years, attributed mainly to the COVID-19 pandemic, where POC tests conducted outside traditional clinical settings played a pivotal role^1,2^. These POC tests are clinical laboratory tests performed close to the patient, give a timely result, and support a clinical decision^3,4^. Various technologies and formats are employed for POC tests, including lateral flow tests, nuclide acid amplification tests, or molecular assays with measured colorimetric, fluorescence, or electrochemical signal properties. In 2019, Land et al. (2019) proposed extending existing POC test characteristics that should be met by defining the acronym REASSURED to catch up with rapid advances in digital and mobile health technology^5^. Accordingly, ideal POC tests fulfill the criteria of **R**eal-time connectivity, **E**ase of specimen collection, **A**ffordable, **S**ensitive, **S**pecific, **U**ser-friendly, **R**apid and robust, **E**quipment-free, and **D**eliverable to end-users^5^.

In the domain of hematology, optical flow cytometry (OFC) stands as the gold standard for quantitative cell analysis with high sensitivity and multiplexing capabilities^6,7^. OFC applications range from HIV diagnostics, cell therapy, virus detection, extracellular vesicle characterization, to cancer prognostics^8,9,10,11,12^. These optical systems comprise lasers, complex optics, and photodetectors with the need for meticulous alignment and calibration^13^. Despite their elaborate technological development, they are primarily restricted to central laboratories with limited access and potential for POC testing. Challenges such as extensive sample pre-treatment, expensive and bulky optics with limited portability, and complex workflow integration hinder compliance with the REASSURED criteria.

To overcome this restraining complexity associated with optical readout systems, magnetic flow cytometers (MFC) with non-optical cell characterization have emerged as an alternative to OFC^14,15,16,17,18,19,20,21,22,23,24,25,26,27,28,29,30^. Utilizing immunologically functionalized magnetic nanoparticles (MNP) instead of fluorescent labels, MFCs employ magnetoresistive sensing mechanisms embedded within microfluidic systems, facilitating miniaturization, cost-effective production, and highly integrated workflows^28^. Due to the non-magnetic nature of biological material, magnetic workflows only require minimal sample pre-processing compared to optical measurement techniques, which encounter issues such as sample matrix opacity and photobleaching^31,32^.

Despite several approaches to develop quantitative cell concentration measurement MFCs, closed-workflow systems that (i) forego sample pre-treatment prior to immunomagnetic labeling and (ii) omit in-between washing or lysis steps or sample transfer to buffer solutions while (iii) enabling on-chip cell quantification are scarcely reported. Shah et al. (2023) employed magnetically labeled agarose microbeads as cell mimics in whole blood (WB) and subsequently detected them with a Hall sensor^29^. Other studies focused on magnetically labeled circulating tumor cells in WB^33,34,35,36,37,38,39^, while Chung et al. (2013) labeled leukocytes magnetically^40^, all followed by magnetic separation but off-chip quantification. In cases where on-chip magnetic cell quantification was performed, cells were manually suspended in a buffer^18,20^, or erythrocytes were manually lysed and sample washed before labeling and measuring to minimize cellular content and remove excess unbound magnetic particles^21^. Reisbeck et al. (2016) labeled erythrocytes and leukocytes in WB, demonstrating that no further sample processing steps are needed for magnetoresistive measurements, while direct cell quantification is feasible^25,27^. Other quantitative approaches focus on bacteria concentration measurements in buffers, including washing steps^22,30^. Overall, the integration of immunomagnetic labeling and quantitative, magnetoresistive sensing, both in WB, poses a significant challenge. Either the labeling process suites magnetic separation methods, or labeling deficiencies need to be remedied by cell lysis and washing steps, albeit at the expense of quantification quality. Conversely, sensor signal analysis suffers from insufficient cell magnetization or false-positive particle aggregates, particularly for micrometer-sized magnetic beads^15,17,30^. Achieving a balance between magnetic particle size, sufficient sensor signal, and appropriate incubation conditions remains challenging in developing closed POC workflow solutions for cell quantification in WB.

In this work, we demonstrate and investigate for the first time holistically concatenated immunomagnetic cell labeling and magnetoresistive sensing, both in WB conditions, including automated signal analysis. Moreover, we established a streamlined MFC workflow for POC testing, encompassing the essential steps (**Figure 1**): sample collection, labeling, sensing, and signal analysis. With a focus on our MFC technology, the sample containing magnetically labeled cells is transported through a microfluidic channel in a laminar flow. Due to a permanent magnetic field, the magnetized cells are attracted to the surface of the chip, initiating a rolling motion. Employing sheathless magnetophoretic enrichment and focusing by magnetic rails, individual magnetized cells are manipulated actuator-free with micrometer precision towards the sensing elements. As these magnetized cells traverse the sensing elements, they induce a change in resistance, while non-magnetic background cells pass through undetected. The distinctive signal signatures generated facilitate multi-parametric cell analysis akin to OFC. To optimize the immunomagnetic cell labeling, we conducted extensive experimental and numerical characterization of MNPs to relate how MNP size influences the magnetic moment of cells. Besides minimizing the MNPs’ induced background noise and false-positive sensor signals, we addressed the temporal effects of immunomagnetic cell labeling regarding bound MNPs, detected cell concentrations, and signal-to-noise ratios in a MFC setup. Finally, we conducted intra-assay benchmarking of the POC workflow under WB conditions with CD14^+^ monocytes, complemented by a robustness assessment across multiple donors to ensure reliability and consistency.

**Figure 1.**
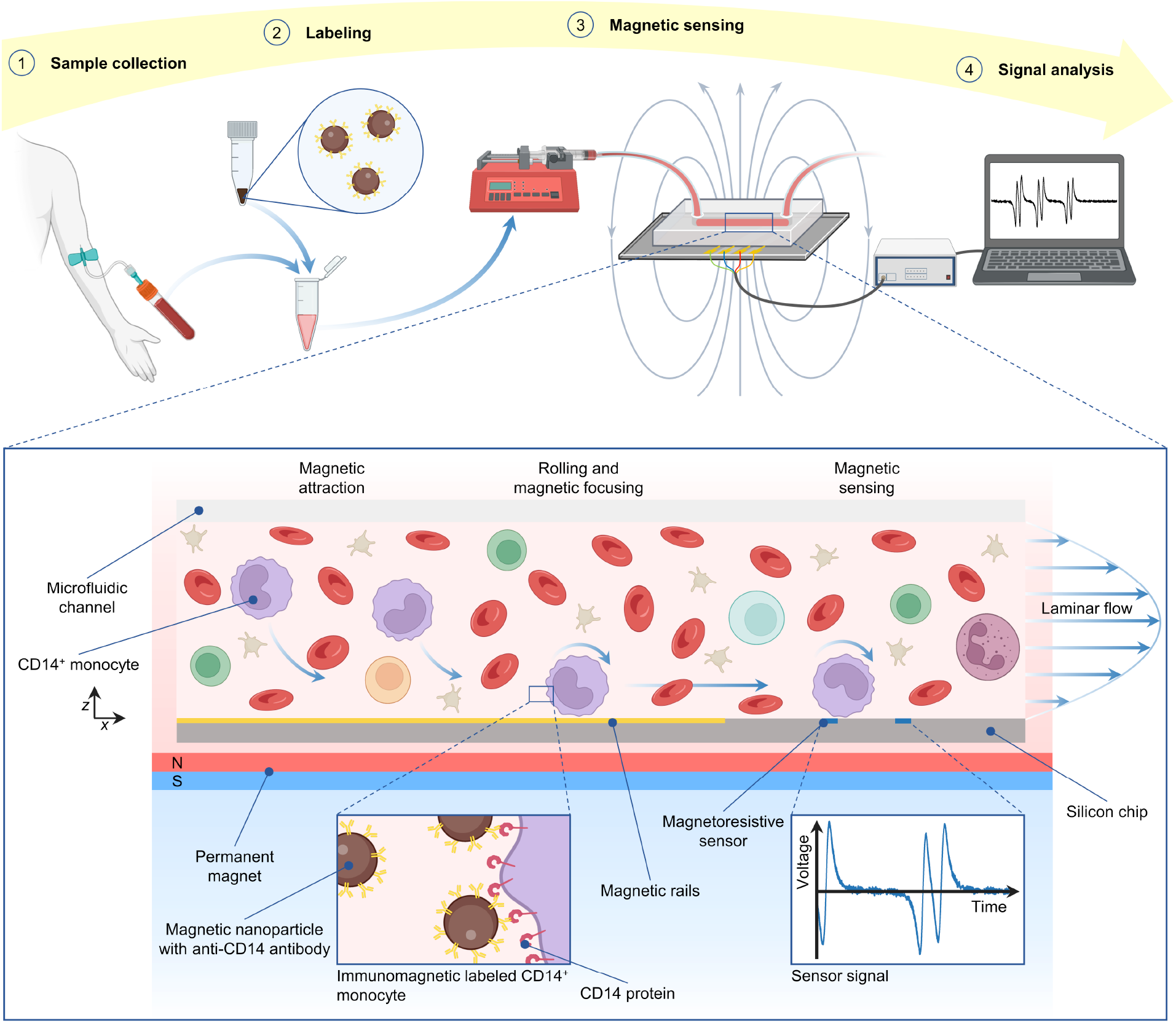
Streamlined magnetic flow cytometry workflow without any washing or cell lysis steps. Collected whole blood is mixed with antibody-coated magnetic nanoparticles to immunomagnetically label target cells, here CD14^+^ monocytes. Without any further processing, the sample is transported through a microfluidic channel, where the magnetically labeled cells are attracted to the silicon (Si) chip surface. Magnetophoretic focusing by magnetic rails precisely focuses the labeled cells on the magnetoresistive sensors. When a labeled cell traverses the sensors, a distinct signal pattern is generated, while the non-magnetic cellular matrix is not detected by the sensors. Automated signal analysis allows for quantitative single cell analysis and cell concentration determination.

## 2. Results and Discussion

### 2.1. Quantitative Magnetic Flow Cytometry in Whole Blood

To establish the MFC’s capability in quantifying cell concentrations in WB, we used magnetic microbeads as cell mimics with known concentrations determined *via* OFC. Across microbead concentrations ranging from 0.15 – 11 µL^-1^ in both Dulbecco’s Phosphate Buffered Saline (DPBS) and WB, the MFC demonstrated linear scalability (R^2^ > 0.99) irrespective of the matrix (**Figure 2** a), b)). The presence of cellular background in WB resulted in a larger 95% confidence interval, indicating that the microbeads traversed over the giant magnetoresistive (GMR) sensors less reliably. In contrast, the concentration deviated for MFC at microbead concentrations above 11 µL^-1^. The coefficient of variation (CV) increased with lower concentrations, with a mean of 0.12 and a standard deviation (SD) of 0.07 for MFC with DPBS, and a mean of 0.13 and an SD of 0.08 in WB, compared to OFC with a mean of 0.06 and an SD of 0.04, combining all concentration measurements (Figure 2 c)). Due to the integration of the magnetic rails and GMR sensors in the bottom of the channel, microbeads must travel maximally the entire channel height (150 µm) and get focused before reaching the sensors. Consequently, a higher cellular background presents more obstacles for the microbeads, counteracting the magnetic attraction and guiding force. As these microbeads traverse the channel longitudinally, they encounter other cells, predominantly erythrocytes, on their way toward the channel bottom. We assume that this interaction prevents some of them from approaching the magnetic rails and GMR sensors closely enough to be focused and detected. Additionally, the cellular background interferes with the microbeads on the magnetic rails, potentially impeding their precise focusing on the GMR sensors. For reference, in 1:20 diluted WB, erythrocyte concentrations are approx. 5 magnitudes higher than the microbead concentration, ranging from 2 – 3 × 10^5^ µL^-1^ ^41^. With the attraction and focusing of the microbeads, an up-concentration in microbeads is achieved. Figure 2 e) shows typical sensor signals for varying numbers of passing microbeads. While the number of microbeads from single- and double-bead events can be reliably inferred, the number of microbeads in clusters of multiple microbeads could not be decoded explicitly. Due to the superposition of the signal from the two Wheatstone half-bridge sensor elements, signal peaks can vanish but not multiply, resulting in an underestimation of the microbead concentration (see double-bead events in Figure 2 e)). Driven by a Poisson distribution, clusters of multiple microbeads passing the sensors are promoted at higher concentrations. Reducing the up-concentration effect by shallower or narrower channels could reduce the cluster formation on the magnetic rails. Compared to the OFC, where the microbeads were suspended in DPBS, the MFC typically detected fewer microbeads (Figure 2 a)). While some errors are unavoidable, consistently occurring errors can be readily rectified. Especially the loss of microbeads is likely in tubing connections and alike, since the magnetophoretic focusing efficiency and detection is expected to be in the range of 98% ^19^. For concentrations below 11 µL^-1^, the error with MFC sizes linearly like the concentration, indicating the optimal concentration range of this setup (Figure 2 a), c), d)).

**Figure 2.**
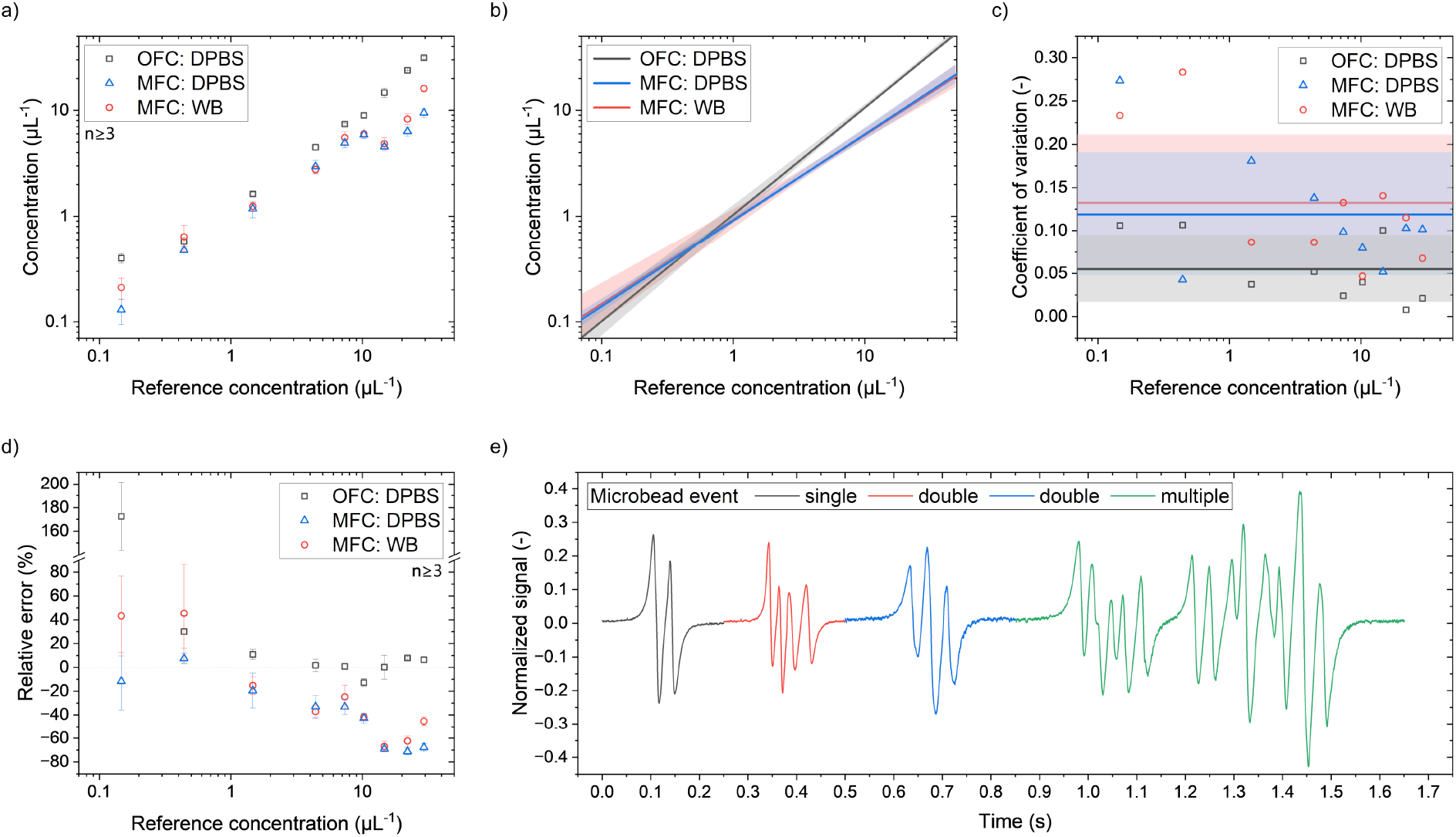
Quantitative magnetic flow cytometer concentration measurements of 12 µm magnetic microbeads in DPBS and 1:20 diluted blood matrix with reference to optical flow cytometer measurements in DPBS. **a)** The concentrations determined with MFC are lower than with OFC and diverge for concentrations above 11 µL^-1^. The blood matrix has little effect on the determined concentration. **b)** Linear fits and 95% confidence intervals for selected concentration intervals. The fits for MFC only include concentrations ≤ 11 µL^-1^, while the fit for OFC includes only concentrations ≥ 0.4 µL^-1^. **c)** The mean over all concentrations of the CV are represented by the horizontal lines with their respective SD. The mean CVs from OFC and MFC measurements in DPBS are 0.06 and 0.12, respectively, while the mean CV from MFC with blood matrix is 0.13. **d)** For concentrations above 0.4 µL^-1^, the relative error with MFC is negative, thus underestimating the concentration in that range. The OFC typically overestimates the concentration. **e)** Typical GMR sensor signals from a varying number of microbeads passing the sensor simultaneously or sufficiently close.

### 2.2. Magnetic Nanoparticle Characteristics

Suppliers offering MNP in kits tailored for specific purposes, typically immunomagnetic cell labeling and separation, usually do not disclose crucial properties of the MNPs, including size, concentration, and magnetic moment. Nevertheless, this information is indispensable when optimizing immunomagnetic labeling processes. Micromod offers MNPs in various sizes, facilitating the general conclusion regarding the dependence of magnetic moment on diameter (**Figure 3** a)). Employing a power law approach, Micromod MNPs can be characterized by the equation m = 2.71678×10^−23^×d^2.98761^, while Chemicell MNPs can be described by m = 9.48583×10^−25^×d2.86401, where m denotes the MNP’s absolute magnetic moment in A m^2^ and d represents the MNP’s diameter in nanometers. Considering that the magnetic moment scales linearly with volume, and a sphere’s volume is a cubic function of its diameter, the similar exponent of the power law fit reflects this relationship, too. However, the magnetic moment can vary several magnitudes among manufacturers within their respective MNP size class, making the highly MNPs from Miltenyi, Stemcell, and Biolegend attractive for MFC applications.

**Figure 3.**
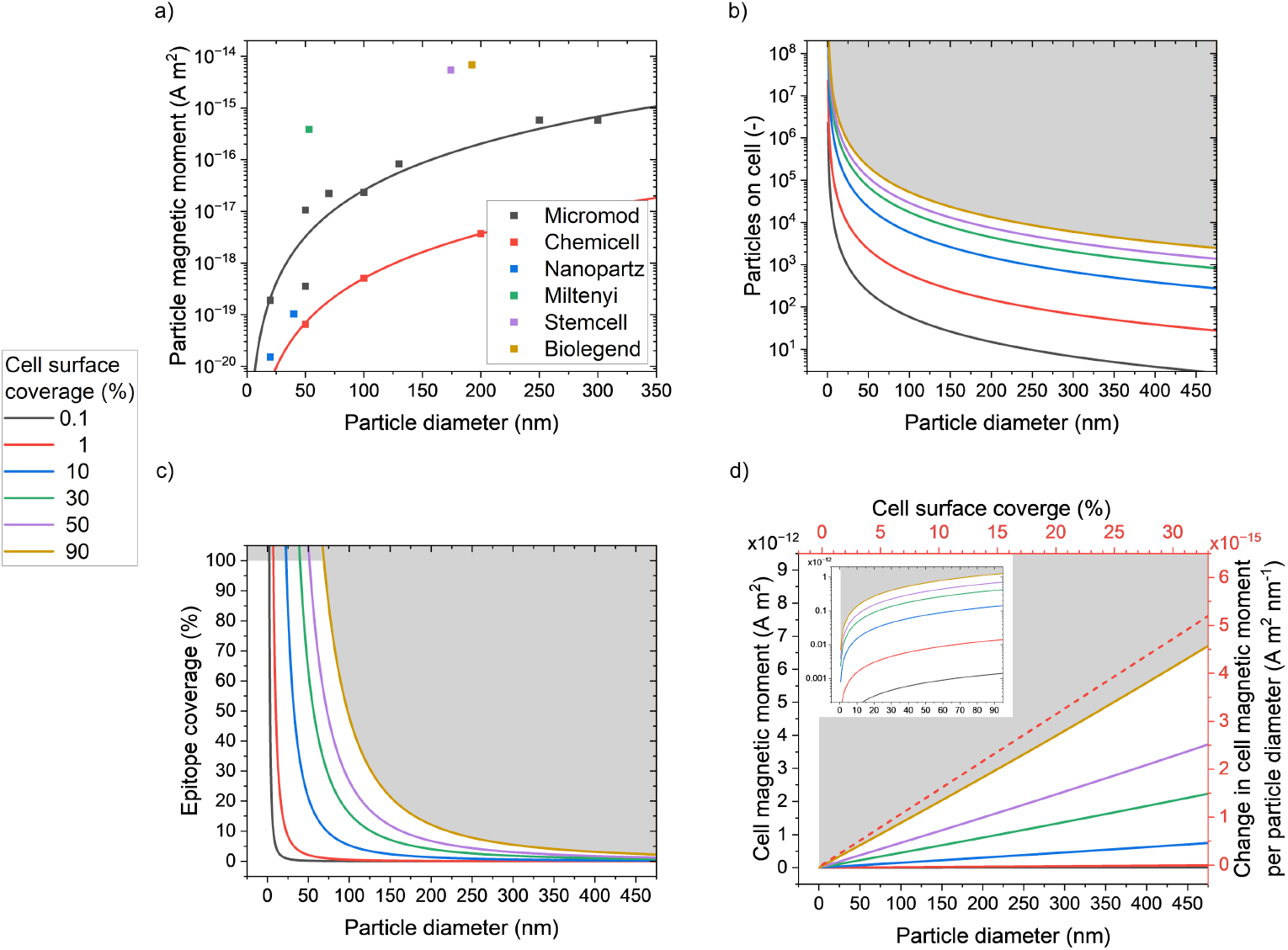
Magnetic nanoparticle characterization and estimation of immunomagnetic labeling results of a 12 µm monocyte. **a)** The particle magnetic moment and diameter dependency follow a power law function. However, different manufacturers achieve different magnetic moments, whereas the particles from commercial immunomagnetic labeling kits (Miltenyi, Stemcell, and Biolegend) show the highest magnetic moments in their particle size class. **b-c)** Assuming no steric hindrance of particles on the cell surface, the number of particles on the cell surface increases exponentially for smaller particles, similar to the epitope coverage limited by the 110,000 CD14 epitopes accessible on the monocyte surface. **d)** Assuming the fitted magnetic moment – diameter relation for micromod particles in combination with the cell coverage by area, the cell magnetization can be estimated based on the particle diameter. Thereby, the magnetic moment increases almost linearly with larger particles. When taking the mean slope of the particle diameter – cell magnetic moment relation for each cell surface coverage, the expected change in cell magnetic moment at a given cell surface coverage when varying the particle size can be estimated (dashed red line with right and top axes), e.g., at 20% cell surface coverage, each nanometer increase in particle diameter would increase the cell magnetic moment by 4.84 × 10^−15^ A m^2^.

The cell surface coverage quantifies the proportion of the cell surface area covered by the total particle’s projected area on the cell’s surface. The theoretical maximum of approx. 90.7% is achieved for a sphere possessing an infinite diameter, thus resembling a plane, while the particle has a finite diameter^42^. While the cell surface coverage in biological systems is rarely quantified, it is anticipated to be a few percent. The relation between particle diameter and particles on a cell scales logarithmically for constant surface coverages (Figure 3 b)). Assuming a monocyte with a diameter of 12 µm and 110,000 CD14 epitopes on its surface, each binding to a particle, the epitope coverage can be determined (Figure 3 c)^43^. Considering the theoretical maximum surface coverage, complete coverage of all epitopes with a single particle can only be achieved with particles smaller than 69 nm. Conversely, 50 nm particles can reach a maximum cell surface coverage of 47% before all epitopes are occupied. To estimate the magnetic moment of labeled cells based on MNP’s diameter, the power law fit for Micromod MNPs is utilized, resulting in a practically linear relationship (Figure 3 d)). By determining the average slopes for different cell surface coverages, the cell magnetic moment and MNP diameter can be combined to describe the expected linear change in the cell magnetic moment for variations in MNP diameter. Essentially, both cell surface coverage and MNP diameter linearly influence the cell magnetic moment. This equal impact on the cell magnetic moment provides two parameters to enhance the immunomagnetic labeling to achieve high cell magnetic moments.

### 2.3. Background Signal Noise from Magnetic Nanoparticles

Implementing washing steps after immunomagnetic labeling to eliminate excess MNPs would disrupt a streamlined workflow. Although unbound MNPs would persist in suspension within the sample, they should not produce any sensor signals or interfere with measurements. Consequently, the volumetric magnetic moment becomes pertinent, with less emphasis on the MNP concentration in the sample, as the MNP size does not primarily determine the quantity of the MNPs bound to a cell. While all MNPs with a volumetric magnetic moment less than 4.20 × 10^−9^ A m^2^ µL^-1^ induced noise lower than the empirically determined limit of 10^−12^ V^2^ in noise signal power, which corresponds to a signal-to-noise ratio (SNR) of ∼7 dB, only MNPs with a diameter of 50 nm and 70 nm fell below the noise limit at the highest volumetric magnetic moment of 1.68 × 10^−8^ A m^2^ µL^-1^ (**Figure 4** a)). However, this observation may not hold for a typical measurement duration of several minutes (Figure 4 c)). Despite the initially low signal noise in the first few seconds of the measurement, signal peaks similar to those of immunomagnetically labeled cells were recorded from MNPs with a diameter of 130 nm after just 1 min, increasing in amplitude and frequency over time. Rarely did MNPs with 70 nm in diameter induce a signal peak protruding from the noise, although the noise was greater than that of MNPs with 50 nm in diameter. Images depicting the magnetic rails reveal the accumulation of MNPs with a diameter of 130 nm, while MNPs with a diameter of 50 nm do not accumulate at the edges of the magnetic rails (Figure 4 d)).

**Figure 4.**
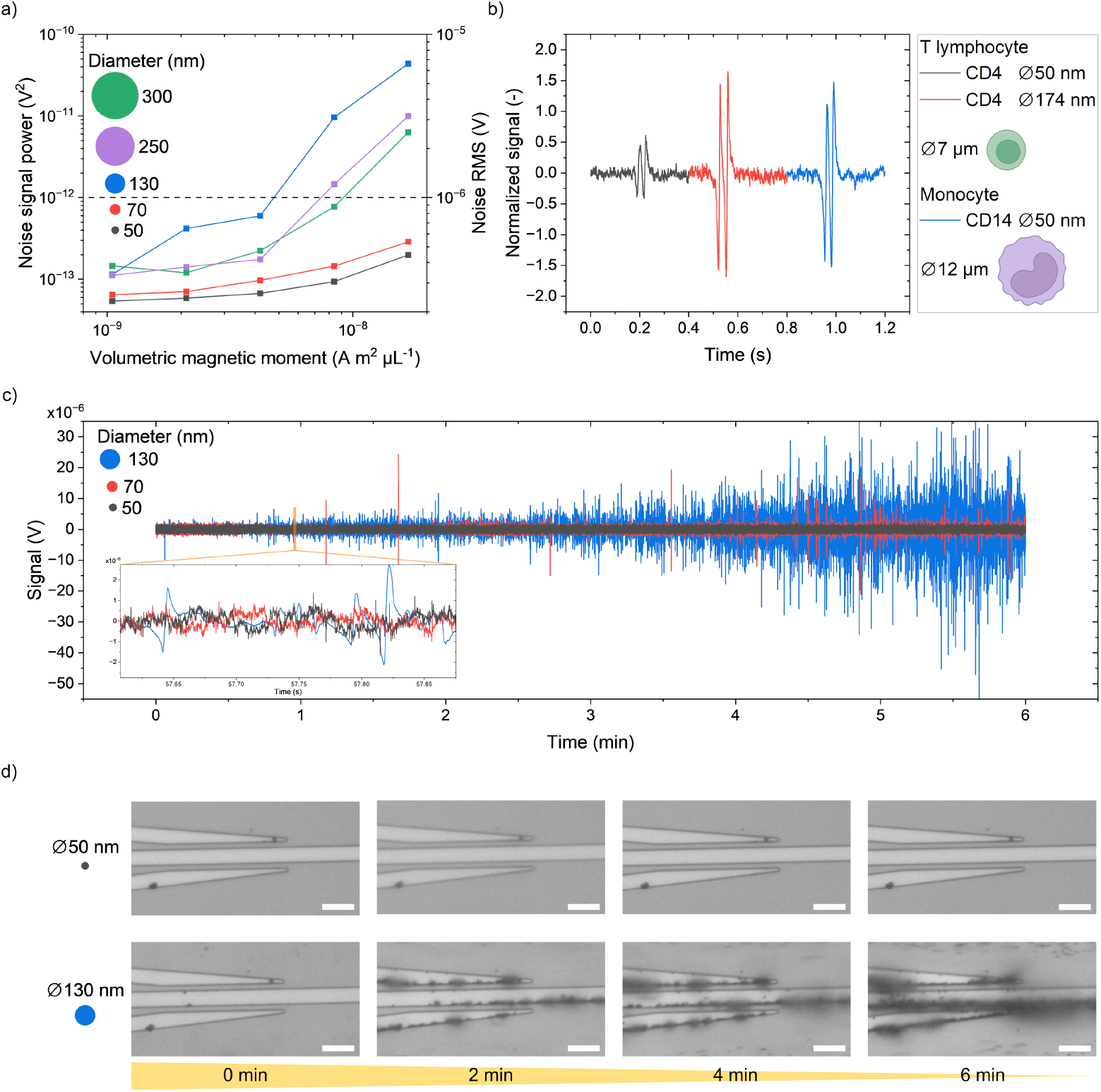
Magnetic nanoparticle background noise. **a)** With a higher volumetric magnetic moment, the sensor noise increases. Additionally, larger particles tend to generate more noise. The noise limit for robust signal analysis is marked with a dashed line at 10^−6^ V. **b)** The particle and cell diameter impact the signal amplitudes. Different particles from commercial labeling kits give different signal amplitudes and noise. Although the particles with 50 nm in diameter are identical except for their antibody functionalization, immunomagnetic labeled CD4^+^ T lymphocytes give less than half the amplitude of CD14^+^ monocytes. **c)** Noise development during the measurement time. After less than 1 min measurement time, 130 nm particles generate signal peaks similar to immunomagnetic labeled cells with further increasing noise. 70 nm particles exceed the noise limit of 10^−6^ V after 1 min. 50 nm particles create no noise larger than the noise limit over the whole measurement period of 6 minutes. **d)** Images from the magnetic rails before the GMR sensors with suspended particles at a volumetric magnetic moment of 1.68 × 10^−8^ Am^2^ µL^-1^ show increasing accumulation of 130 nm magnetic nanoparticles, whereas 50 nm magnetic nanoparticles do not agglomerate at the magnetic rails. The scale bars represent 20 µm.

Although the MNPs themselves would not generate signals akin to those from immunomagnetically labeled cells, their aggregation could produce a sufficient magnetic moment for sensor detection. The elevated magnetic field gradients at the magnetic rail borders focus immunomagnetically labeled cells but also encourage MNP aggregation. When these aggregates reach a critical size, the laminar flow force detaches them, transporting them across the sensors, and the aggregate induces a sensor signal. To mitigate false-positive sensor signal occurrences, MNPs with diameters of 50 nm or less are most suitable.

### 2.4. Comparing the Immunomagnetic Labeling of Different Cell and Magnetic Nanoparticle Sizes

MFC is based on detecting alterations in a magnetic field induced by passing magnetized cells, which exhibit a linear correlation with the signal from the GMR sensor. Although various cells are expected to yield comparable MNP cell surface coverages, their potentiality to evoke magnetic field changes by their magnetic fringe field hinges on the absolute number of bound MNPs and, consequently, their cell size. The average fringe field acting on the GMR sensor’s free layer, generated by a sufficiently homogeneously labeled cell, can be approximated using a magnetic dipole field:

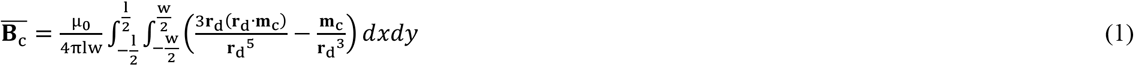

where µ_0_ is the magnetic permeability in vacuum, l is the GMR sensor length, w is the GMR sensor width, **r**_d_ is the dipole’s position vector, and **m**_**c**_ = **Σ m**_**i**_ is the cell’s magnetic moment derived from the sum of individual MNP magnetic moments **m**_i_. While the magnetic fringe field scales linearly with the cell’s magnetic moment, the position vector exerts an inversely cubic impact 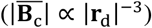. During cell rolling over the GMR sensor, the minimal position vector corresponds to the cell’s radius 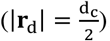. Hence, the cell diameter is expected to dominate over the cell surface and epitope coverage in generating high signal amplitudes. For instance, a CD4^+^ T lymphocyte, with a diameter of 7 µm and ∼46,000 CD4^+^ epitopes, is 1.7 times smaller and possesses 2.4 times fewer epitopes for MNPs to bind than a CD14^+^ monocyte^44^. With a similar MNP cell surface coverage, the magnetic dipole field of a CD4^+^ T lymphocyte is expected to be ∼2-fold stronger than that of a CD14^+^ monocyte, resulting in a similar higher GMR signal amplitude.

However, when using MNPs with a 50 nm diameter and the exact magnetic moment to label CD4^+^ T lymphocytes and CD14^+^ monocytes, the latter yields an almost threefold higher signal amplitude or magnetic dipole field (Fig 4 b)). The SNR for the CD4^+^ T lymphocyte is 8.26 dB while the CD14^+^ monocyte has an SNR of 17.71 dB. This discrepancy can only be rationalized by a substantially lower cell surface coverage with MNPs of the CD4^+^ T lymphocyte, indicative of inadequate and non-reproducible labeling conditions. Only by substituting the 50 nm MNPs with 174 nm diameter MNPs having a 14-fold higher magnetic moment for CD4^+^ T lymphocyte labeling a similar signal amplitude (or an SNR of 18.9 dB) to that of the CD14^+^ monocyte could be attained (Fig 4 b)). This implies that ∼25.2 times fewer MNPs were bound to the CD4^+^ T lymphocyte, with an MNP diameter favoring false-positive signals from MNP aggregates. Albeit the MNPs were designed explicitly for immunomagnetic cell separation from WB, their suitability for MFC applications needs to be individually reviewed.

### 2.5. Immunomagnetic Cell Labeling Characteristics in Whole Blood

The dynamics of immunomagnetic cell labeling were assessed based on the signal amplitudes of the GMR sensors and the number of detected cells using MFC relative to OFC. In addition to MNP size and magnetic moment, the incubation time and antibody binding affinity in WB influence the quantity of MNPs bound to the cell, as reflected by their magnetic moment and sensor signal amplitude. Despite the manufacturer’s recommended incubation time of 15 min for the 50 nm MNPs, signal amplitudes generated after 15 – 30 min were either insufficient or nonexistent for signal analysis. Investigating the effect of incubation time on signal amplitude from immunomagnetically labeled CD14^+^ monocytes revealed that only after approx. 1.5 h were signal amplitudes of 7 × 10^−6^ V sufficiently high enough for reliable detection (**Figure 5** a)). Incubation times ranging from 1.5 – 4 h did not significantly increase the signal amplitudes from CD14^+^ monocytes. However, an extended incubation period led to a second plateau in signal amplitudes at around 26 × 10^−6^ V after 5.5 – 6 h. The observed signal response aligns well with a biphasic dose-response function (R^2^ > 0.91) (see parameters in Supplement Section 2). While antibody-diffusion and -binding reactions typically transact within minutes, the subsequent increase in signal amplitude after 4.5 h suggests MNP aggregation on MNPs already bound to CD14^+^ monocytes. Investigation into cell labeling dynamics within the first hour of incubation was not feasible due to insufficient signal amplitude. Incubating at room temperature compared to the recommended 2 – 8 °C may be one reason for the change in antibody performance.

**Figure 5.**
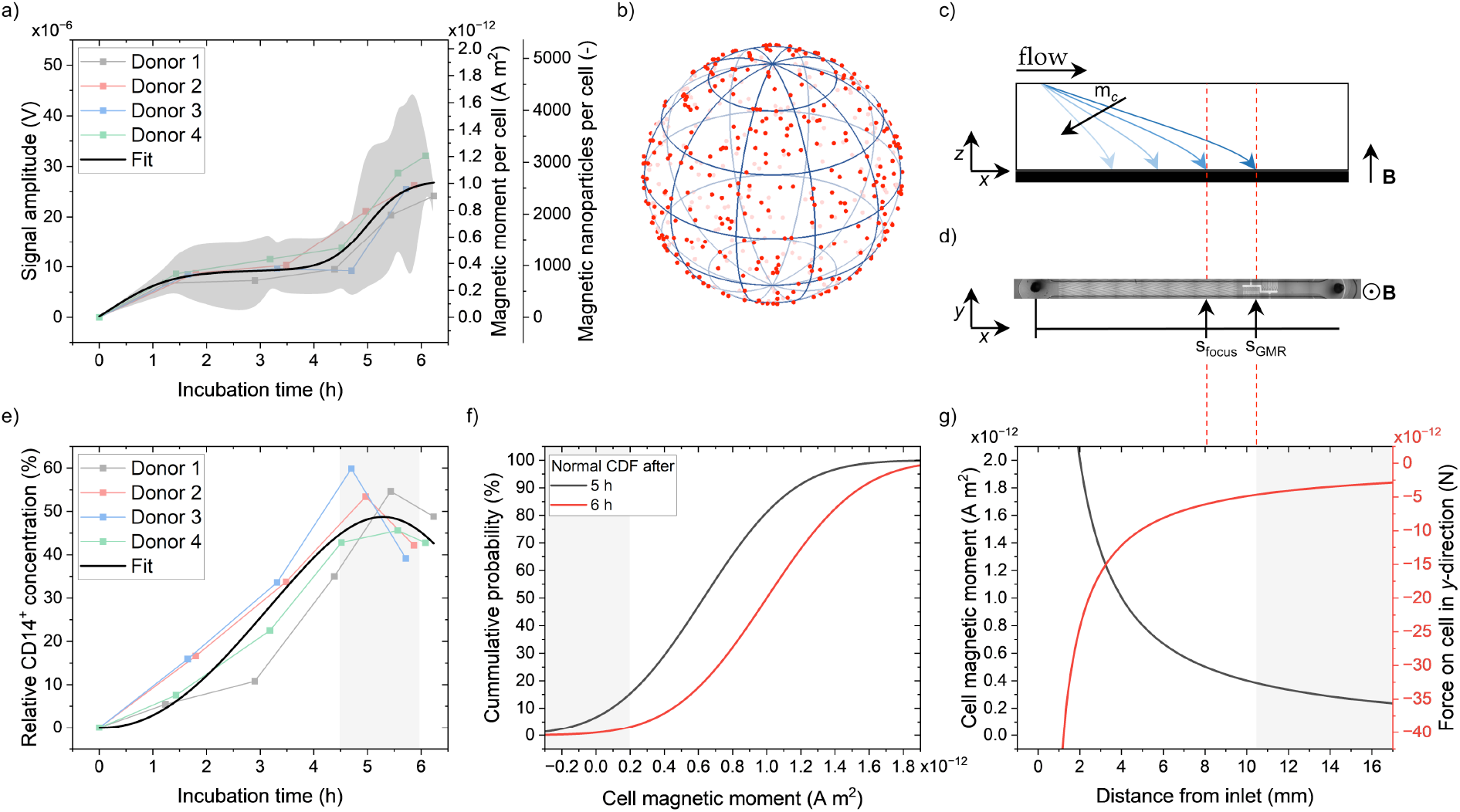
Immunomagnetic labeling of CD14^+^ monocytes in whole blood and quantification with magnetic flow cytometry. **a)** The signal amplitude increases with longer incubation, forming a plateau between 1.5 h and 4 h. The grey area displays the SD. **b)** Model of a labeled monocyte with 500 MNPs (red) homogeneously distributed on its surface. The MNPs are displayed five-fold larger in the case of an MNP diameter of 50 nm. **g)** Depending on the magnetic moment of the cell, a force is exerted on the cell in *y*-direction. At a flow rate Q of 20 µL min^-1^, the cell reaches the Si chip when traveling the channel height of 150 µm at a distance from the inlet, depending on its magnetic moment. The grey area marks the distances after the location of the GMR sensors (s_*x*_ ≥ s_GMR_ ≈ 10.5 mm). **c)** Trajectories of attracted cells depending on their magnetic moment m_c_ in a side view perspective. **d)** Top view image from sensor chip with magnetic rails and GMR sensor array. s_GMR_ and s_focus_ denote the distance from the inlet to the GMR sensors and the last magnetic rail element, respectively. **e)** The relative concentration with respect to the optical reference increases with longer incubation times, peaks after approx. 5 h, and tends to decrease for even more prolonged incubation. **f)** Normal cumulative probability density functions (CDF) for the cell magnetic moment after an incubation period of 5 h and 6 h. The gray area marks the cell’s magnetic moments below the detection limit.

Utilizing an MFC, the signal amplitude is related to the cell’s magnetic moment, assuming the magnetized cell behaves as a magnetic dipole. To address the relationship between the cell’s diameter and its magnetic moment, Reisbeck et al. (2016) conducted numerical simulations of GMR sensor signals, resulting in calibration lines for various cell diameters^27^. Here, the change in magnetic moment relative to the signal amplitude is established as a general model to amalgamate these discrete calibration lines into a single calibration curve. Neglecting the initial magnetization of MNPs **M**_**0**_ due to a saturated magnetization of the MNPs in a strong magnetic flux density (B_ext_ > 100 mT), the magnetic dipole moment **m** can be given as

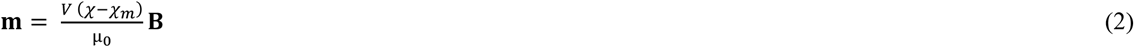

where **B** represents the magnetic flux density, χ is the MNPs’ volume susceptibility, χ_m_ is the matrix’s volume susceptibility, V denotes the MNPs volume, and µ_0_ is the vacuum magnetic permeability^45,46,47^. Given that the cell surface coverage remains constant across different cell sizes, the volume of the MNPs scales with the cell volume, thus a cubic function is deemed suitable for fitting the absolute magnetic moment of a cell (R^2^ > 0.997):

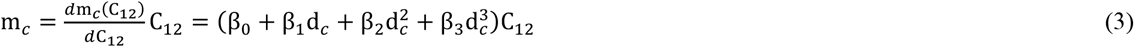

Here, C_12_ denotes the average peak-to-peak signal amplitude, and d_c_ is the cell diameter. With

β_0_ = 1.42092 × 10^−11^ A m^2^ V^-1^,

β_1_ = 3.13750 × 10^−10^ A m^2^ V^-1^,

β_2_ = 4.38506 × 10^−11^ A m^2^ V^-1^,

β_3_ = 4.63497 × 10^−12^ A m^2^ V^-1^,

C_12_ in V, and d_c_ in micrometers, Equation (3) yields values in A m^2^. The cell diameter can be inferred from the peak-normalized integral of the characteristic four-peak signal (see Supplement Section 3)^27^. Regardless of the incubation time, a mean hydrodynamic diameter of CD14^+^ monocytes of 12.2 µm and a SD of 1.7 µm were determined. With this model, the least detectable magnetic moment of a CD14^+^ monocyte using our MFC is estimated to be approx. 2 × 10^−13^ A m^2^. Considering the known magnetic moment of a single MNP, a minimum number of > 500 bound MNPs per cell is required for detection as a magnetized cell, assuming a CD14^+^ monocyte diameter of 12.2 µm. After incubating ∼6 h, an average of around 2500 MNPs were bound on a CD14^+^ monocyte, which corresponds to a cell surface coverage of approx. 0.2%.

### 2.6. How Cell Quantification Depends on Immunomagnetic Labeling

In contrast to the development of the signal amplitude, the relative CD14^+^ monocyte concentration does not exhibit an intermediate plateau but peaks after approx. 5 h (Figure 5 e)). Within the initial 4.5 h, more MNPs are not necessarily bound to a detected cell; instead, more cells surpass the minimal signal amplitude threshold by binding more MNPs for detection. Approx. 50% of the CD14^+^ monocytes, compared to OFC, were detected at most before the concentration decreased after 5.5 h. This decline could be attributed to CD14^+^ monocyte death, although binding kinetics may also play a role, where dissociation of antibody bonds dominates over association kinetics.

The magnetic detection of only half of the CD14^+^ monocytes could also be attributed to insufficient magnetic moments to be attracted by the magnetic field to the sensor chip. Additionally, the high cellular background presents many obstacles that must be circumvented when traversing the maximum distance of the channel height (150 µm) before passing the sensor. The primary forces governing the attraction of magnetized cells to the sensor chip surface are the magnetic force **F**_**m**_ and fluidic drag force **F**_**d**_, expressed as:

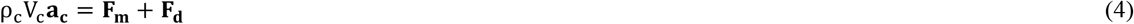

where ρ_c_ and V_c_ denote the density and volume of the cell, including bound MNPs, respectively, and **a**_**c**_ represents the cell’s acceleration. A magnetic flux density gradient and the magnetic moment of the labeled cell **m**_**c**_ that is modeled as a magnetic dipole generate the magnetic force:^47,48,49^

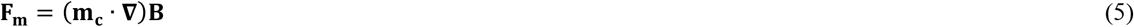

The fluidic drag force partially counteracts the magnetic force and arises from the relative movement of the cell to the suspending matrix:

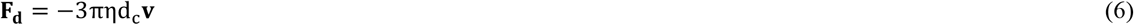

where η is the dynamic viscosity of the suspending matrix, d_c_ is the labeled cell’s hydrodynamic diameter, and **v** is the relative velocity of the labeled cell to the matrix. From the position and size of our permanent magnet, we can suppose the magnetic flux density components only in the *z*-direction, neglecting all the others, and a saturated magnetic moment at the applied magnetic flux density of ∼150 mT. In our MFC, the relevant direction of cell attraction is towards the sensor chip in the *z*-direction. To calculate the time t_*z*_ for a labeled cell to travel the distance s_*z*_, we employ Equation (4) – (6), neglecting the inertia of the labeled cell, and use **v = s t**^**-1**^:

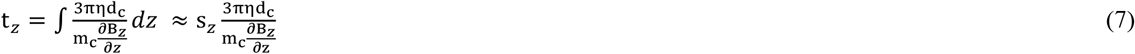

The approximation on the right side of Equation (7) is only valid for a constant magnetic flux density gradient over the distance of s_*z*_. During this traveling time t_*z*_, the labeled cell is transported longitudinally in the laminar sample flow in the *x*-direction, s_*x*_, and must reach the sensor chip surface before passing the GMR sensors at position s_GMR_. When a labeled cell needs to travel the complete channel height h (s_*z*_ = h), the transported distance by the laminar flow s_*x*_ can be determined using the mean flow velocity 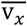:

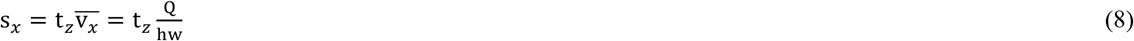

where Q is the sample flow rate, and w is the channel width.

Inserting Equation (7) into Equation (8), and considering the viscosity of water (η = 8.9 × 10^−4^ Pa s), the flow rate Q of 20 µL min^-1^, the cell diameter d_c_ of 12.2 µm, the channel dimensions (s_z_ = h = 150 µm, w = 700 µm), and the experimentally determined magnetic flux density gradient 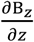 of –12.13 T m^-1^ (see Supplement Section 5), the minimal magnetic moment of the cell and respective force required to be timely on the sensor surface for proper detection by the GMR sensors can be determined (Figure 5 g)). Since the GMR sensors are located 10.5 mm from the inlet, a minimum cell magnetic moment of 3.8 × 10^−13^ A m^2^ is needed for detection (Figure 5 d), g). However, for reliable focusing of labeled cells onto the GMR sensors by the magnetic rails, the labeled cell must reach the sensor chip within 6.5 mm from the inlet, necessitating a cell magnetic moment of 6.3 × 10^−13^ A m^2^. On average, this threshold is achieved after approx. 5 h of incubation, although significant variability persists (Figure 5 a)).

In Figure 5 f), normal cumulative distribution functions (CDF) for incubation times of 5 h and 6 h are depicted with mean magnetic moments of 6.3 × 10^−13^ A m^2^ and 10 × 10^−13^ A m^2^ and typical SDs of 4.2 × 10^−13^ A m^2^ and 3.5 × 10^−13^ A m^2^, respectively. Given the MFC’s sensor detection limit of approx. 2 × 10^−13^ A m^2^, approx. 15% of the labeled cells will remain undetected after 5 h of incubation and reduce to 3% after 6 h, resembling the loss from insufficient labeling. Considering that approx. 50% of the CD14^+^ monocytes were detected overall, the remaining loss ranging between 35% – 45% likely stems from MFC setup itself and the influence of the cellular background, which slightly surpasses losses observed with the reference microbeads (see Section 2.1).

### 2.7. CD14^+^ Monocyte Quantification with Streamlined Magnetic Flow Cytometry Workflow

While the statistical cell representation with approx. 10% is limited after a 1.5 h incubation period, we opted to incubate between 4.5 – 6.5 h in favor of a higher cell detection yield, consequently agreeing less with POC testing demands (Figure 5 b)). To assess the reproducibility of our streamlined MFC workflow, we processed 4 blood sample tubes from a single donor through identical steps and conducted 3 measurements for each tube **(Figure 6** a)). The CV within sample triplicates (precision) was at most 0.15, while the overall intra-assay CV did not surpass 0.11. Replicates from 4 different donors yielded a maximum CV of 0.14 (Figure 6 b). On average, the relative concentration to OFC was 50% with an SD of 0.05, resulting in an overall CV of 0.10. This bias can be probably well compensated to improve accuracy since linearity is provided over a sufficient concentration range. Despite the limited number of statistical samples tested, our streamlined MFC workflow exhibits precision and robustness with CVs comparable to a typical regulatory approval threshold of ≤ 0.20 ^50^.

**Figure 6.**
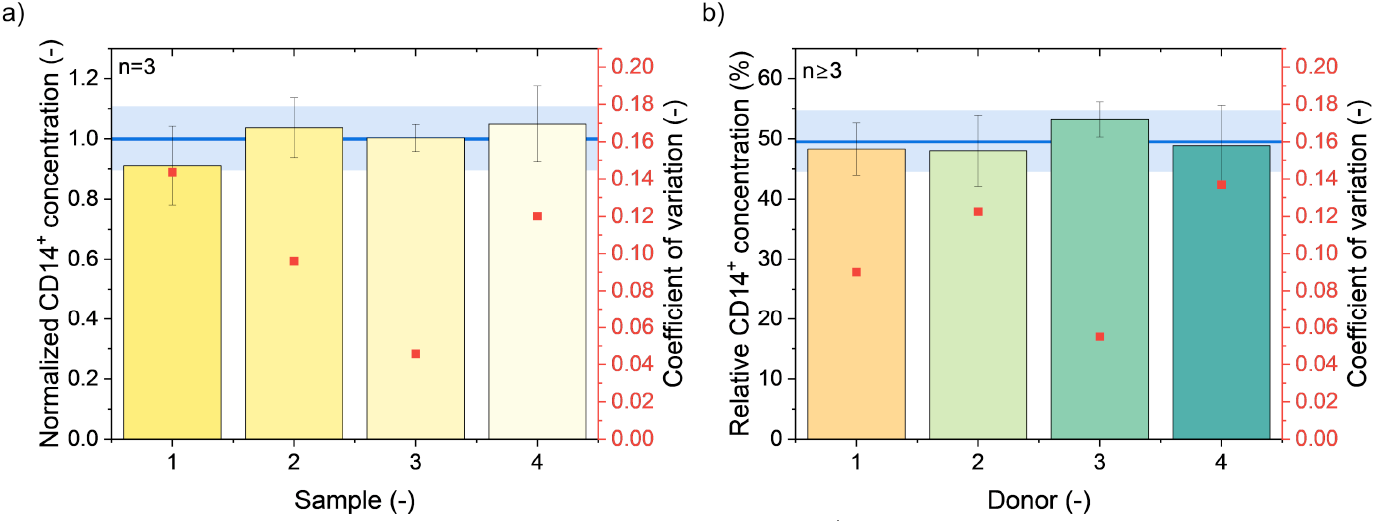
The streamlined MFC workflow performance with CD14^+^ monocytes in whole blood. **a)** Triplicates from 4 collecting tubes drawn from the same donor with incubation times between 5 – 6 h have CV values lower than 0.15. The blue line and bar indicate the overall mean and SD, resulting in a CV of 0.11. **b)** Blood from four different donors was measured with incubation times between 4.5 – 6.5 h. The MFC’s determined concentration is roughly half the optical reference’s concentration. The blue line and bar indicate the overall mean and SD, resulting in a CV of 0.10.

## 3. Conclusions

MFC has excellent potential for POC testing applications but lacks integrated workflow solutions that allow on-chip cell labeling and quantification. Typically, erythrocyte lysis precedes immunomagnetic labeling to improve the labeling quality, or external washing steps remove cellular background and MNP excess to allow subsequent cell quantification in a buffer medium, at which MFC workflows aiming for POC testing are essentially interrupted. This demonstrated that MFC can quantify cell concentrations in WB without any washing or cell lysis steps. Our streamlined workflow only involved the steps of sample collection, immunomagnetic cell labeling, magnetoresistive detection, and signal analysis.

We first used reference microbeads as a biomimetic model to demonstrate linearity in cell concentration quantification in WB over a dynamic range of almost two logarithmic orders, limited only by inaccurate signal analysis at concentrations > 11 µL^-1^. The cellular background from WB slightly increased the variance for concentrations < 1 µL^-1^. Our MFC showed high up-concentration potential, with the concentration range adjustable by adapting the channel geometry, making it suitable for quantifying rare cells like circulating tumor cells.

Furthermore, we investigated the characteristics of MNPs and their role in immunomagnetic labeling. Our findings include a correlation between MNP size, magnetic moment, and the resulting implications for the final cell magnetic moment. Omitting washing steps left unbound MNPs in the sample, contributing to background signal noise. Depending on the MNP concentration, size, and measurement time, noise induced by MNPs could reach levels of labeled cell signals, including cell-like signals from MNP agglomerates. Although larger MNPs typically result in higher cell magnetic moments, their background noise can easily lead to false-positive signals, making them unsuitable for cell quantification without washing steps. We concluded that MNPs with diameters < 50 nm are most suitable to minimize the risk of false-positive signals while typically providing sufficient magnetic moment.

Moreover, we discussed the dynamics of immunomagnetic cell labeling in WB, focusing on incubation time and resulting magnetic moment. Our results with CD14^+^ monocytes indicate that long incubation periods are necessary for sufficient cell labeling and quantification with minimal background. Theoretical considerations about sensor signal and cell magnetic moment allowed us to quantify the bound MNPs per cell, revealing insights into labeling quality. We were able to detect CD14^+^ monocytes with 12.2 µm in diameter having bound only 500 MNPs, equivalent to a magnetic moment of approx. 2 × 10^−13^ A m^2^. Comparing different cell and MNP diameters, we experience cell- and MNP-specific labeling qualities, favoring smaller cells for MFC due to the inversely cubic relation between the cell magnetic fringe field inducing the GMR sensor signal and their diameter 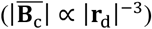. However, the hours-long incubation times might infringe on POC testing requirements. Antibody binding kinetics remain imperfect for MFC applications and need to be reviewed individually, although they were initially designed for labeling and purifying cells in WB.

Additionally, we explored how cell quantification depends on immunomagnetic labeling and factors influencing cell detection efficiency by MFC with integrated sample preparation. While the magnetic rails and GMR sensors integrated in the bottom of the channel allow actuator-free focusing and sensing, two conditions need to be met for reliable detection: (i) Labeled cells need to be attracted onto the sensor chip and focused on the GMR sensors, which depends on the chosen flow rate, cell size and magnetic moment. (ii) Cells must have sufficient magnetic moment to be detected by the GMR sensors. Consequently, a higher cellular background could impede cell movement toward the sensors, resulting in lower detection efficiency. Longer incubation times lead to higher detection rates, whereas our findings suggest that a remaining error results from failing to comply with the first condition. This interaction with other cells, mainly erythrocytes, and insufficient magnetic moment hinders some CD14^+^ monocytes from approaching the sensors closely enough to be detected.

Finally, we benchmarked our streamlined MFC workflow in WB conditions with CD14^+^ monocytes. Without needing cell lysis or washing steps, we showed an intra-assay CV of 0.11 across four samples. We demonstrated the robustness of the workflow with four donors, reaching a CV of 0.10, comparable to regulatory approval thresholds.

Future efforts to optimize MFC for POC testing should address elaborate signal analysis strategies to increase the dynamic concentration range next to the MNP design. Approaches to implementing artificial intelligence for time series analysis seem to be a promising tool to entangle the signal from overlapping microbead events^51^. With a focus on tailored antibodies for WB applications with fast reaction kinetics and mechanical stress resistance, cell magnetic moment and reliable cell quantification could be significantly improved.

Conclusively, our study provides valuable insights into optimizing and applying MFC for quantitative cell analysis in complex biological matrices like WB, underscoring its potential as a reliable tool for quantitative cell analysis in clinical settings, particularly in POC testing scenarios.

## 4. Experimental Section

### 4.1. Size, Concentration and Magnetic Moment of Nanoparticles

The concentration and size of the nanoparticles from Chemicell (chemicell GmbH, Berlin, Germany), Micromod (micromod Partikeltechnologie GmbH, Rostock, Germany), and Nanopartz (Nanopartz Inc., Loveland, USA) were taken from their respective data sheets. For particles from Biolegend (BioLegend, San Diego, USA) Miltenyi (Miltenyi Biotech B.V. & Co. KG, Bergisch Gladbach, Germany) and Stemcell (STEMCELL Technologies Canada Inc., Vancouver, Canada), the nanoparticle size was determined with Multi-Angle Dynamic Light Scattering (MADLS) in a Zetasizer Ultra Red Label (Malvern Panalytical Ltd., Worcestershire, United Kingdom) from at least six measurements at 25 °C. Magnetite (Fe_3_O_4_) with a refractive index of 2.36 and absorption of 0.147 was used for the material model. The nanoparticles were diluted in ultrapure Milli-Q water to a suitable concentration, and finally, the size was determined by volume. The concentration was obtained with the same device from the average of at least 9 measurements with nanoparticles diluted in different concentrations in Milli-Q water or sterile filtered Dulbecco’s Phosphate Buffered Saline (DPBS) (Sigma-Aldrich Corp., St. Louis, USA).

The average particle magnetic moment was obtained from vibrating sample magnetometer (VSM) (PMC MicroMag 3900-CF, Lake Shore Cryotronics Inc., Westerville, USA) measurements. 20 µL of the respective particle solution in stock concentration was pipetted in the center of rolled-up 2 × 1 cm^2^ cellulose paper, which was beforehand inserted into empty gelatin capsules of 5 mm in diameter and 16 mm in length. After drying, the non-magnetic capsules were closed. For measurement, a capsule was mounted in a custom-made sample holder for the VSM, the vibration amplitude set to 1 mm and frequency to 83 Hz. In the following, a hysteresis loop was recorded by varying the magnetic field between ± 300 mT in increments of 5 mT and measuring the magnetic moment of the sample averaged over 200 ms. At saturation, the magnetic moment can be read from the hysteresis plot. However, a slope correction was performed beforehand to correct for ferromagnetic components that do not show saturation. The final average particle magnetic moment is obtained by dividing the magnetic moment by the nanoparticle concentration. Exemplary hysteresis plots can be found in the Supplement Section 1.

**Table 1** gives an overview of all investigated nanoparticles. Measured properties are marked with [*]; all others are taken from the respective data sheets.

**Table 1.**
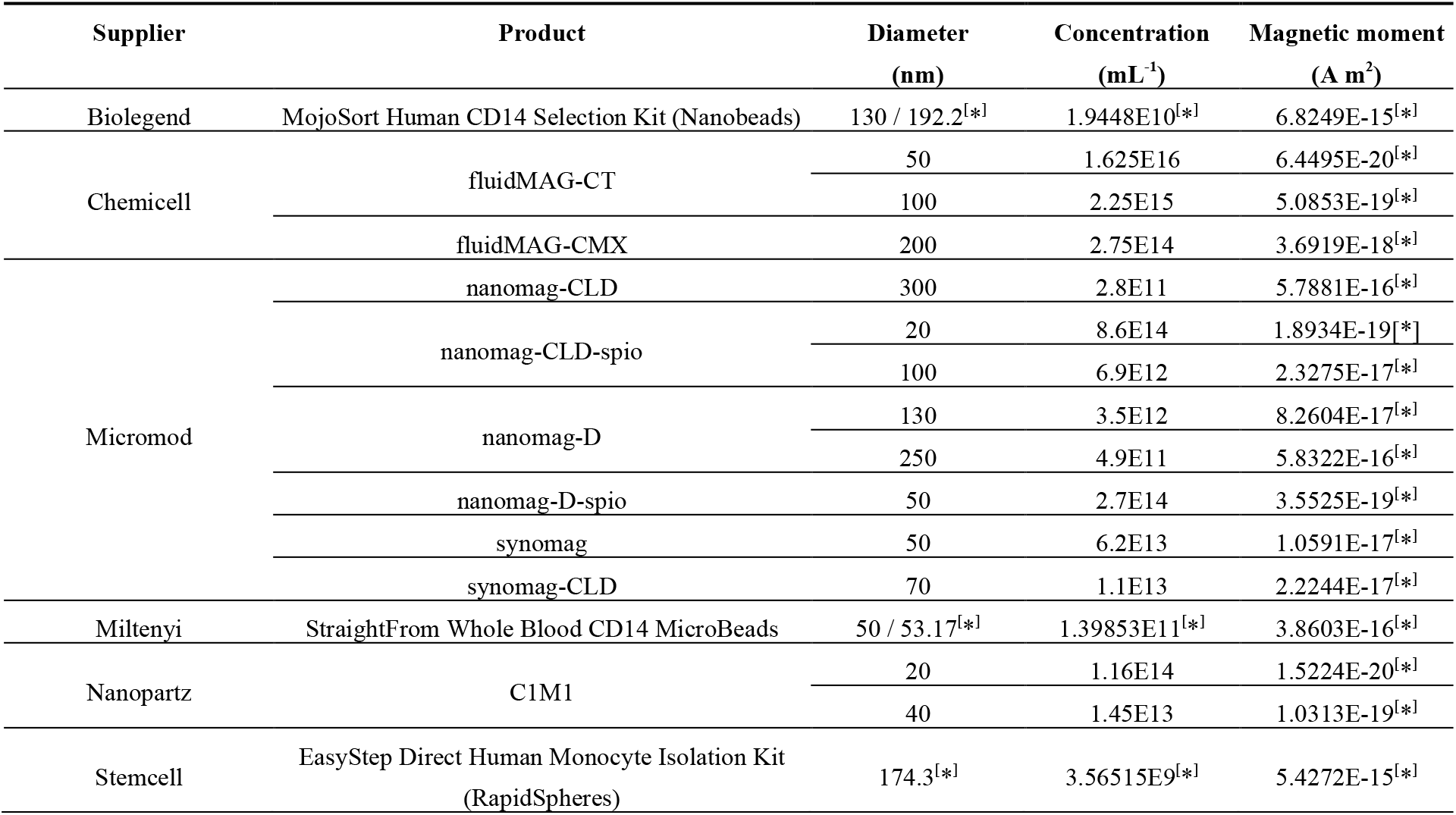
Size, concentration and magnetic moment of nanoparticles.

### 4.2. Composition of the Magnetic Flow Cytometer

The core of the magnetic flow cytometer is the structured silicon (Si) chip of 1 × 2 cm^2^ in size comprising giant magnetoresistive (GMR) sensing elements and magnetic rails that were fabricated by Sensitec GmbH (Wetzlar, Germany). The GMR sensors had a width of 2 µm and a length of 30 µm configured in a Wheatstone half-bridge configuration. The distances between two sensing elements within a bridge varied from 14 – 18 µm, modulating the signal length. Magnetic rails were arranged in chevron-like patterns on the Si chip consisting of a 150 nm chromium bottom layer and a 200 nm nickel iron top layer. These rails facilitated the precise magnetic focusing on the GMR sensors of magnetized objects on the Si chip. A 70 nm thick silicon nitride layer covered the whole Si chip, except the electrical connection pads, to protect the GMR sensors, magnetic rails, and electrical lines from corrosive blood. The Si chip was mounted on a printed circuit board (PCB), and both were electrically connected by wire bonding. Beneath the Si chip, a NdFeB permanent magnet of 32 × 27 × 6 mm^3^ in size (NE3227, IBS Magnet, Berlin, Germany) was precisely positioned to generate an external magnetic field density B_ext_ of ∼150 mT in the sensor plane with a magnetic field density gradient maximally perpendicular to the GMR sensors. This external magnetic field density gradient was used to attract the magnetized cells or magnetic microbeads toward the Si chip and as a bias field for the GMR sensors.

On top of the Si chip, a straight fluidic channel made of polydimethylsiloxane (PDMS) was positioned over the magnetic rails and GMR sensors. Standard soft-photolithography processes were used to fabricate a negative master mold of epoxy resin (SU-8 2050, Kayaku Advanced Materials Inc., Westborough, USA) on a 4” Si wafer substrate as described elsewhere^52^. Inlet and outlet ports were punched into the PDMS with a biopsy punch (WellTech Rapid Core 0.5 mm ID, World Precision Instruments LLC, Sarasota, USA) before assembly. Subsequently, the fluidic was stamped on a thin layer of uncured PDMS spun on a Si wafer before aligning the channel onto the Si chip under a stereo microscope. Baking the assembly at 60 °C for 2 h cures the PDMS, forming a stable bond between the fluidic and Si chip. Inserting polytetrafluoroethylene (PTFE) tubing (RCT-ZS-DKA-SW 0.3 mm inner diameter, RCT Reichelt Chemietechnik GmbH + Co., Heidelberg, Germany) into the outlet and inlet, and connecting the latter via a dosing tip (0.013” inner diameter, Nordson Corp., Westlake, USA) with a glass syringe (1750 TLL, Hamilton Company Corp., Reno, USA) completed the fluidics. Pulsation-free sample flow was facilitated by a syringe pump (Fusion 4000, Chemyx Inc., Stafford, USA)^53^.

During a measurement, the Wheatstone half-bridge was supplied with an AC modulated signal (MFLI 500 kHz Lock-in Amplifier, Zurich Instruments AG, Zurich, Switzerland) with a frequency of 10 kHz and a peak-to-peak voltage of 0.9 V or 1 V. The differential signal of the Wheatstone half-bridge was filtered with a 3^rd^ order low-pass filter having a time constant of 300 µs and amplified 5k or 10k-fold by a lock-in amplifier (MFLI 500 kHz Lock-in Amplifier, Zurich Instruments AG) before being digitized by a data acquisition board (NI USB-6351, National Instruments Corp., Austin, USA) with a sample rate of 10 kS s^-1^ and a resolution of 16 bit and recorded with a custom written PC program (LabVIEW 2018, National Instruments Corp.). A customized microscope with 20× magnification and equipped with a CMOS camera system (GS3-U3-32S4M-C, Teledyne FLIR LLC, Wilsonville, USA) was used for optical control of the measurement conditions.

Since the GMR sensor sensitivity, supply voltage, and signal amplification directly scale the recorded signal, the signal stream was normalized with the respective values. Subsequently, the characteristic signal pattern was detected and counted with a custom algorithm implemented in a finite-state machine as described elsewhere^27^.

### 4.3. Reference Microbeads

To demonstrate quantitative concentration measurements with the magnetic flow cytometer, we mimicked cells with polystyrene microbeads coated with a superparamagnetic iron oxide shell and 12 µm in diameter (micromer-M, micromod Partikeltechnologie GmbH). These microbeads have an average magnetic moment of 1.01 × 10^−11^ A m^2^ per microbead and were diluted with DPBS and 0.1% polysorbate 20 (Tween 20, Sigma-Aldrich Corp.) or spiked into 1:20 DPBS-diluted whole blood. Microbead dilutions were performed from a single stock containing approx. 147 microbeads per µL, which were finally measured with the OFC (MACSQuant Analyzer 10, Miltenyi Biotech B.V. & Co. KG) or the MFC at a flow rate of 25 µL min^-1^ for 2 min.

### 4.4. Determining Background Noise from Magnetic Nanoparticles

The background signal noise from unbound nanoparticles at different concentrations was quantified and compared with nanoparticle stock solutions diluted in DPBS to have the exact magnetic moment per volume. A sample was flushed for 90 s into the channel at 40 µL min^-1^, then 60 s at 20 µL min^-1^, with an ensuing signal recording for 30 s. After signal normalization, the root-mean-square (RMS) signal noise and power were determined from the 30 s signal stream with a custom-written PC script (MATLAB R2020b, The MathWorks Inc., Natick, USA).

For the recording of the signal development over time, the external magnetic field was initially removed, the channel flushed with nanoparticles diluted in DPBS to a magnetic moment concentration of 1.68 × 10^−8^ A m^2^ µL^-1^, and finally, the external magnetic field reapplied. The subsequent recording was performed with a flow rate of 20 µL min^-1^ and image acquisition every second minute.

The signal-to-noise ratios (SNR) were calculated from the average signal amplitudes A_signal_ and 3-times the noise’s SD: SNR_db_ = 10 *log*_10_ A_signal_^2^ − 10 *log*_10_ 3^2^SD_noise_^2^.

### 4.5. Immunomagnetic Cell Labeling and Magnetic Flow Cytometry Measurements

Fresh venous blood was collected from healthy donors and stabilized with EDTA (S-Monovette EDTA K3E 9mL, Sarstedt AG & Co. KG, Nümbrecht, Germany). For immunomagnetic labeling of CD14^+^ monocytes, a commercial labeling kit for monocyte extraction from whole blood was used, consisting of magnetic nanoparticles functionalized with anti-CD14 antibodies (StraightFrom Whole Blood CD14 MicroBeads, Miltenyi Biotech B.V. & Co. KG). Two parts of whole blood were mixed with one part of magnetic nanoparticles in an Eppendorf tube and incubated in a shaker at room temperature for 1.25 – 6.25 h, depending on the experiment. Finally, the opaque sample was diluted at a ratio of 1:20 or 1:30 with DPBS to a concentration between 1 – 5 µL^-1^ matching the MFC’s dynamic range and measured with a flow rate of 20 µL min^-1^ or 25 µL min^-1^. Neither washing nor lyses steps were performed at any point in the workflow.

Immunomagnetic labeling of CD4^+^ T lymphocytes was performed with two commercial labeling kits. (i) Dextran-coated MNPs and anti-CD4 antibodies (EasySep Human CD4 Positive Selection Kit II, STEMCELL Technologies Canada Inc.) were incubated at a 1:1 ratio over night at 3 – 8 °C. The immunologically functionalized particles were then incubated with WB in a 1:9 ratio for 30 min in a rotation shaker. (ii) MNPs coated with anti-CD4 antibodies (StraightFrom Whole Blood CD4 MicroBeads, Miltenyi Biotech B.V. & Co. KG) were directly mixed with WB in a 1:2 ratio and incubated for 5 – 6 h (analogous to CD14^+^ monocytes) in a rotation shaker.

### 4.6. Optical Reference Measurements

Optical reference measurements to determine the CD14^+^ monocyte concentration were performed with an optical flow cytometer (MACSQuant Analyzer 10, Miltenyi Biotech B.V. & Co. KG). Therefore, 20 µL WB was stained with 1 µL each of anti-CD14 antibody conjugated with FITC fluorophore (REAfinity, Miltenyi Biotech B.V. & Co. KG) and anti-CD45 antibody conjugated with VioBlue fluorophore (REAfinity, Miltenyi Biotech B.V. & Co. KG). The sample was diluted at a ratio of 1:30 (MAQSQuant Running Buffer, Miltenyi Biotech B.V. & Co. KG) after an incubation time of 30 min and directly measured.

## CRediT authorship contribution statement

**Moritz Leuthner**: Conceptualization, Methodology, Software, Validation, Formal Analysis, Investigation, Data Curation, Writing – Original Draft, Writing – Review & Editing, Visualization. **Michael Helou**: Conceptualization, Methodology, Validation, Formal Analysis, Investigation, Resources, Writing – Review & Editing, Funding Acquisition. **Mathias Reisbeck**: Conceptualization, Methodology, Writing – Review & Editing, Funding Acquisition. **Oliver Hayden**: Conceptualization, Resources, Writing – Review & Editing, Supervision, Project Administration, Funding Acquisition.

## Data availability

The data supporting the conclusions of this article will be made available upon reasonable request.

## Acknowledgments

The authors would like to acknowledge Leopold Daum (Technical University of Munich) for sharing his expertise in immunomagnetic labeling and MFC. Furthermore, the authors would like to thank Anika Kwiatkowski (Technical University of Munich) for her support with wire bonding, Robin Karl (Technical University of Munich) for providing access to the Zetasizer, and Michael Wack (Ludwig-Maximilians-Universität München) for facilitating the use of their vibrating sample magnetometer.

This study (406/20 S-EB) was approved by Ethikkomission an der Technischen Hochschule München, approved on 19 October 2020. Informed consent was obtained from each study participant.

This research was funded by the Federal Ministry for Economic Affairs and Climate Action (BMWK) on the basis of a decision by the German Bundestag, grant number ZF4792002AP9.

## Supplementary data

Supplementary data to this article can be found online.

## References

1. Karako, K., Song, P., Chen, Y., Tang, W. Increasing demand for point-of-care testing and the potential to incorporate the Internet of medical things in an integrated health management system. BioScience Trends 16, 4–6 (2022).

2. Valera, E., Jankelow, A., Lim, J., Kindratenko, V., Ganguli, A., White, K., Kumar, J., Bashir, R. COVID-19 Point-of-Care Diagnostics: Present and Future. ACS Nano 15, 7899–7906 (2021).

3. Drain, P.K., Hyle, E.P., Noubary, F., Freedberg, K.A., Wilson, D., Bishai, W., Rodriguez, W., Bassett, I.V. Evaluating Diagnostic Point-of-Care Tests in Resource-Limited Settings. Lancet Infect. Dis. 14, 239–249 (2014).

4. World Health Organization, 2023. Laboratory and point-of-care diagnostic testing for sexually transmitted infections, including HIV (2023).

5. Land, K.J., Boeras, D.I., Chen, X.-S., Ramsay, A.R., Peeling, R.W. REASSURED diagnostics to inform disease control strategies, strengthen health systems and improve patient outcomes. Nature Microbiology 4, 46–54 (2019).

6. De Rosa, S.C., Brenchley, J.M., Roederer, M. Beyond six colors: A new era in flow cytometry. Nat. Med. 9, 112–117 (2003).

7. Perfetto, S.P., Chattopadhyay, P.K., Roederer, M. Seventeen-colour flow cytometry: unravelling the immune system. Nature Immunology 4, 648–655 (2004).

8. Brussaard, C.P.D.; Marie, D., Bratbak, G. Flow cytometric detection of viruses. J. Virological Methods 85, 175–182 (2000).

9. Demaret, J., Varlet, P., Trauet, J., Beauvais, D., Grossemy, A., Hégo, F., Yakoub-Agha, I., Labalette, M. Monitoring CAR T-cells using flow cytometry. Cytometry 100, 115–253 (2021).

10. Douek, D.C., Brenchley, J.M., Betts, M.R., Ambrozak, D.R., Hill, B., Okamoto, Y., Casazza, J.P., Kuruppu, J., Kunstman, K., Wolinsky, S., Grossman, Z., Dybul, M., Oxenius, A., Price, D.A., Connors, M., Koup, R.A. HIV preferentially infects HIV-specific CD+ T cells. Nature 417, 95–98 (2002).

11. Malcovati, L., Germing, U., Kuendgen, A., Della Porta, M.G., Pascutto, C., Invernizzi, R., Giagounidis, A., Hildebrandt, B., Bernasconi, P., Knipp, S., Strupp, C., Lazzarino, M., Aul, C., Cazzola, M. Time-Dependent Prognostic Scoring System for Predicting Survival and Leukemic Evolution in Myelodysplastic Syndromes. J. of Clinical Oncology 25, 3503–3510 (2007).

12. Shen, W., Guo, K., Adkins, G.B., Jiang, Q., Liu, Y., Sedano, S., Duan, Y., Yan, W., Wang, S.E., Bergersen, K., Worth, D., Wilson, E.H., Zhong, W. A Single Extracellular Vesicle (EV) Flow Cytometry Approach to Reveal EV Heterogeneity. Angew. Chem. Int. Ed. 57, 15675–15680 (2018).

13. Dorn-Beineke, A., Sack, U. Quality control and validation in flow cytometry. J. Lab. Med. 40, 1–13 (2016).

14. Boser, B.E., Murali, P. Flow Cytometer-on-a-Chip. IEEE Biomedical Circuits and Systems Conference (BioCAS) Proceedings, 2014, 480–483 (2014).

15. Chícharo, A., Martins, M., Barnsley, L.C., Taouallah, A., Fernandes, J., Silva, B.F.B., Cardoso, S., Diéguez, L., Espiña, B., Freitas, P.P. Enhanced magnetic microcytometer with 3D flow focusing for cell enumeration. Lab Chip 18, 2593–2603 (2018).

16. Duarte, C.M., Fernandes, A.C., Cardoso, F.A., Bexiga, R., Cardoso, S.F., Freitas, P.J.P. Magnetic Counter for Group B Streptococci Detection in Milk. IEEE Trans. Magn. 51, 5100304 (2015).

17. Fernandes, A.C., Duarte, C.M., Cardoso, F.A., Bexiga, R., Cardoso, S., Freitas, P.P. Lan-on-Chip Cytometry Based on Magnetoresistive Sensors for Bacteria Detection in Milk. Sensors 14, 15496–15524 (2014).

18. Giraud, M., Delapierre, F.-D., Wijkhuisen, A., Bonville, P., Thévenin, M., Cannies, G., Plaisance, M., Paul, E., Ezan, E., Simon, S., Fermon, C., Féraudet-Tarisse, C., Jasmin-Lebras, G. Evaluation of In-Flow Magnetoresistive Chip Cell-Counter as a Diagnostic Tool. Biosensors 9, 105 (2019).

19. Helou, M., Reisbeck, M., Tedde, S.F., Richter, L., Bär, L., Bosch, J.J., Stauber, R.H., Quandt, E., Hayden, O. Time-of-flight magnetic flow cytometry in whole blood with integrated sample preparation. Lab Chip 13, 1035–1038 (2013).

20. Huang, C.-C., Ray, P., Chan, M., Zhou, X., Hall, D.A. An aptamer-based magnetic flow cytometer using matched filtering. Biosensors and Bioelectronics 169, 112362 (2020).

21. Issadore, D., Chung, J., Shao, H., Liong, M., Ghazani, A.A., Castro, C.M., Weissleder, R., Lee, H. Ultrasensitive Clinical Enumeration of Rare Cells ex Vivo Using a Micro-Hall Detector. Sci. Transl. Med. 4, 141ra92 (2012).

22. Issadore, D., Chung, H.J., Chung, J., Budin, G., Weissleder, R., Lee, H. µHall Chip for Sensitive Detection of Bacteria. Adv. Healthcare Mater. 2, 1224–1228 (2013).

23. Kokkinis, G., Cardoso, S., Keplinger, F., Giouroudi, I. Microfluidic platform with integrated GMR sensors for quantification of cancer cells. Sensors and Actuators B: Chemical 241, 438–445 (2017).

24. Lee, C.-P., Lai, M.-F., Huang, H.-T., Lin, C.-W., Wie, Z.-H. Wheatstone bridge giant-magnetoresistance based cell counter. Biosensors and Bioelectronics 57, 48–53 (2014).

25. Leuthner, M., Reisbeck, M., Helou, M., Hayden, O. Towards a Point-of-Care Test of CD4+ T Lymphocyte Concentrations for Immune Status Monitoring with Magnetic Flow Cytometry. Micromachines 15, 520 (2024).

26. Murali, P., Niknejad, A.M., Boser, B.E. CMOS Microflow Cytometer for Magnetic Label Detection and Classification. IEEE J. Solid-State Circuits 52, 543–555 (2017).

27. Reisbeck, M., Helou, M.J., Richter, L., Kappes, B., Friedrich, O., Hayden, O. Magnetic fingerprints of rolling cells for quantitative flow cytometry in whole blood. Sci. Rep. 6, 32838 (2016).

28. Reisbeck, M., Richter, L., Helou, M.J., Arlinghaus, S., Anton, B., van Dommelen, I., Nitzsche, M., Baßler, M., Kappes, B., Freidrich, O., Hayden, O. Hybrid integration of scalable mechanical and magnetophoretic focusing for magnetic flow cytometry. Biosensors and Bioelectronics 109, 98–108 (2018).

29. Shah, N., Iyer, V., Zhang, Z., Gao, Z., Park, J., Yelleswarapu, V., Aflatouni, F., Johnson, A.T.C., Issadore, D. Highly stable integration of graphene Hall sensors on a microfluidic platform for magnetic sensing in whole blood. Microsystems & Nanoengineering 9, 71 (2023).

30. Soares, A.R., Afonso, R., Martins, V.C., Palos, C., Pereira, P., Caetano, D.M., Carta, D., Cardoso, S. On-site magnetic screening tool for rapid detection of hospital bacterial infections: Clinical study with Klebsiella pneumonia cells. Biosensors and Bioelectronics: X 11, 100149 (2022).

31. Gaster, R.S., Hall, D.A., Nielsen, C.H., Osterfeld, S.J., Yu, H., Mach, K.E., Wilson, R.J., Murmann, B., Liao, J.C., Gambhir, S.S., Wang, S.X. Matrix-insensitive protein assays push the limits of biosensors in medicine. Nature Medicine 15, 1327–1332 (2009).

32. Issadore, D., Par, Y.I., Shao, H., Lee, K., Liong, M., Weissleder, R., Lee, H. Magnetic sensing technology for molecular analysis. Lab Chip 14, 2385–2397 (2014).

33. Gribko, A., Stiefel, J., Liebetanz, L., Nagel, S.M., Künzel, J., Wandrey, M., Hagemann, J., Stauber, R.H., Freese, C., Gül, D. IsoMAG – An Automated System for the Immunomagnetic Isolation of Squamous Cell Carcinoma-Derived Circulating Tumor Cells. Diagnostics 11, 2040 (2021).

34. Hoshino, K., Huang, Y.-Y., Lane, N., Huebschman, M., Uhr, J.W., Frenkel, E.P., Zhang, X. Microchip-based immunomagnetic detection of circulating tumor cells. Lab Chip 11, 3449–3457 (2011).

35. Kim, S., Han, S.-I., Park, M.-J., Jeon, C.-W., Joo, Y.D., Choi, I.-H., Han, K.-H. Circulating Tumor Cell Microseparator Based on Lateral Magnetophoresis and Immunomagnetic Nanobeads. Anal. Chem. 85, 2779–2786 (2013).

36. Kirby, D., Glynn, M., Kijanka, G., Ducrée, J. Rapid and Cost-Efficient Enumeration of Rare Cancer Cells from Whole Blood by Low-Loss Centrifugo-Magnetophoretic Purification Under Stopped-Flow Conditions. Cytometry Part A 87A, 74–80 (2015).

37. Liu, W., Nie, L., Li, F., Aguilar, Z.P., Xu, H., Xiong, Y., Fu, F., Xu, H. Folic acid conjugated magnetic iron oxide nanoparticles for nondestructive separation and detection of ovarian cancer cells from whole blood. Biomater. Sci. 4, 159–166 (2016).

38. Nie, L., Li, F., Huang, X., Aguilar, Z.P., Wang, Y.A., Xiong, Y., Fu, F., Xu, H. Folic Acid Targeting for Efficient Isolation and Detection of Ovarian Cancer CTCs from Human Whole Blood Based on Two-Step Binding Strategy. ACS Appl. Mater. Interfaces 10, 14055–14062 (2018).

39. Xu, H., Aguilar, Z.P., Yang, L., Kuang, M., Duan, H., Xiong, Y., Wei, H., Wang, A. Antibody conjugated magnetic iron oxide nanoparticles for cancer cell separation in fresh whole blood. Biomaterials 32, 9758–9765 (2011).

40. Chung, J., Issadore, D., Ullal, A., Lee, K., Weissleder, R., Lee, H. Rare cell isolation and profiling on a hybrid magnetic/size-sorting chip. Biomicrofluidics 7, 054107 (2013).

41. Rosen, R. Optimality principles in biology (Butterwoth & Co (Publishers) Ltd., London, 1967).

42. Clare, B.W., Kepert, D.L. The closest Packing of Equal Circles on a Sphere. Proc. R. Soc. Lond. A 405, 329–344 (1986).

43. Antal-Szalmas, P., Van Strijp, J.A.G., Weersink, A.J.L., Verhoef, J., Van Kessel, K.P.M. Quantitation of surface CD14 on human monocytes and neutrophils. J. of Leukocyte Biology 61, 721–728 (1997).

44. Poncelet, P., Poinas, G., Corbeau, P., Devaux, C., Tubiana, N., Muloko, N., Tamalet, C., Chermann, J.C., Kourilsky, F., Sampol, J. Surface CD4 density remains constant on lymphocytes of HIV-infected patients in the progression of disease. Res. Immunol. 142, 291–298 (1991).

45. Gijs, M.A.M. Magnetic bead handling on-chip: new opportunities for analytical applications. Microfluid. Nanofluid. 1, 22–40 (2004).

46. Liu, C., Stakenborg, T., Peeters, S., Lagae, L. Cell manipulation with magnetic particles toward microfluidic cytometry. J. of Applied Physics 105, 102014 (2009).

47. Shevkoplyas, S.S., Siegel, A.C., Westervelt, R.M., Prentiss, M.G., Whitesides, G.M. The force acting on a superparamagnetic bead due to an applied magnetic field. Lab Chip 7, 1294–1302 (2007).

48. Gassner, A.-L., Abonnc, M., Chen, H.-X., Morandini, J., Josserand, J., Rossier, J.S., Busnel, J.-M., Girault, H.H. Magnetic forces produced by rectangular permanent magnets in static microsystems. Lab Chip 9, 2356–2363 (2009).

49. Zborowski, M., Sun, L., Moore, L.R., Williams, P.S., Chalmers, J.J. Continuous cell separation using novel magnetic quadrupole flow sorter. J Magnetism and Magnetic Materials 194, 224–230 (1999).

50. European Medicines Agency, Committee for Medical Products for Human Use, 2022. ICH guideline M10 on bioanalytical method validation and study sample analysis, European Medicines Agency, The Netherlands, available online: https://www.ema.europa.eu/en/documents/scientific-guideline/ich-guideline-m10-bioanalytical-method-validation-step-5_en.pdf (accessed on 17 April 2024) (2022).

51. Baur, M., Reisbeck, M., Hayden, O., Utschick, W. Joint Particle Detection and Analysis by a CNN and Adaptive Norm Minimization Approach. IEEE Transactions on biomedical engineering 69, 2468–2479 (2022).

52. Xia, Y., Whitesides, G.M. Soft Lithography. Angew. Chem. Int. Ed. 37, 550–575 (1998).

53. Leuthner, M., Hayden, O. Grease the gears: how lubrication of syringe pumps impacts microfluidic flow precision. Lab Chip 24, 56–62 (2024).

