## Supplementary Information for "Quantitative Magnetic Flow Cytometry in High Hematocrit Conditions for Point-of-Care Testing"

M. Helou
EarlyBio GmbH, Bottroper Weg 2, 13507 Berlin, Germany

### 1. VSM hysteresis curves of MNPs

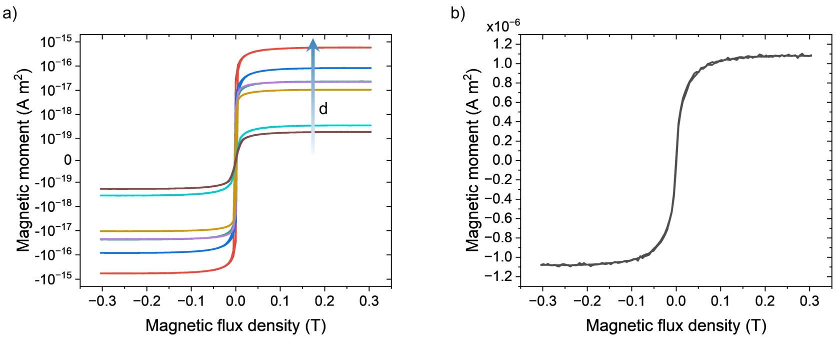


**Figure S1.** Hysteresis curves that are scaled for a single particle from Micromod MNPs (**a**)) for increasing diameters d and from the Miltenyi MNPs with 50 nm in diameter (**b**)). The magnetic moment is saturated for all particles at the MFC's operation conditions of 150 mT.

### 2. Fitting function for signal amplitude

A biphasic dose response function was used to fit the signal amplitudes:

$$A(t) = A_1 + (A_2 - A_1)\left[\frac{p}{1+10^{(L_1-t)h_1}} + \frac{1-p}{1+10^{(L_2-t)h_2}}\right] \quad \text{(S1)}$$

with

$A_1 = -6.26505 \times 10^{-12}$

$A_2 = 2.7318 \times 10^{-5}$

$L_1 = 0.00819$

$L_1 = 0.20795$

$h_1 = 18.20023$

$h_1 = 29.42125$

$p = 0.46262$

and time t in h gives amplitude A(t) in V.

#### 3. Determination of the hydrodynamic cell diameter from the signal integral

The hydrodynamic cell diameter $d_c$ can be inferred with a cubic fit of the peak-normalized distance integral $I_{xna}$ of the four-peak GMR signal:

$$d_c(I_{xna}) = \beta_0 + \beta_1 I_{xna} + \beta_2 {I_{xna}}^2 + \beta_3 {I_{xna}}^3 \qquad \text{(S2)}$$

with

$\beta_0 = -1.30763$

$\beta_1 = 0.51154$

$\beta_2 = -0.00909$

$\beta_3 = 3.7124 \times 10^{-4}$

The parameters $\beta_0$ to $\beta_3$ were obtained from numerical signal simulations (Reisbeck et al., 2016). The peak-normalized distance integral is given by:

$$I_{xna} = v_c \sum_{i=1}^{4} \int \frac{|A_i(t)|}{\max|A_i(t)|} dt \qquad \text{(S3)}$$

where $v_c$ denotes the cell velocity determined from the peak and GMR sensor distances, and $A_i(t)$ denotes the amplitude signal from the $i^{th}$ peak.

#### 4. Determination of the cell magnetic moment from the signal amplitude

To determine the cell magnetic moment, Reisbeck et al. (2016) numerically simulated a calibration line connecting sensor signal amplitude with magnetic moment for each cell diameter. The slope of each calibration line can be extracted. Fitting these slopes with a cubic function relates the change in magnetic moment with respect to the signal amplitude to the cell's diameter (Figure S2). The red values from the numerical data were omitted, since they were affected by numerical instabilities. The resulting fitted curve has an adjusted $R^2$ of 0.9995:

$$\frac{d\mathrm{m}_c(\mathrm{C}_{12})}{d\mathrm{C}_{12}} = \beta_0 + \beta_1 \mathrm{d_c} + \beta_2 \mathrm{d_c}^2 + \beta_3 \mathrm{d_c}^3 \qquad \text{(S4)}$$

with

$\beta_0 = 1.42092 \times 10^{-11}$

$\beta_1 = 3.13750 \times 10^{-10}$

$\beta_2 = 4.38506 \times 10^{-11}$

$\beta_3 = 4.63497 \times 10^{-12}$

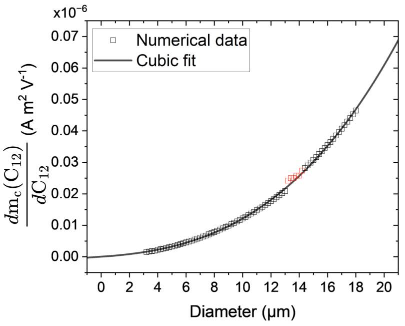


**Figure S2.** General model for inferring the cell magnetic moment from the signal amplitude with respect to the hydrodynamic cell diameter. Values in red are omitted for the cubic fit.

### 5. Determination of the magnetic flux density gradient

The magnetic flux density is measured with a Gauss meter (CYHT201, Chenyang Technologies GmbH & Co. KG, Finsing, Germany) in 250 µm steps moving perpendicular from the magnet (Figure S3). With a cubic fit ($R^2$ > 0.999) and linear interpolation starting at a magnetic flux density of 150 mT in the sensor plane and using the channel height of 150 µm, the magnetic flux density gradient along the channel height $\frac{\partial B_z}{\partial z}$ is determined to – 12.13 T m$^{-1}$. The analytical model for the magnetic flux density of a permanent magnet is determined by (Gou et al., 2004):

$$B_{ext}(z) = \frac{B_r}{\pi}\left[arctan\left(\frac{lw}{2z\sqrt{4z^2+l^2+w^2}}\right) - arctan\left(\frac{lw}{2(h+z)\sqrt{4(h+z)^2+l^2+w^2}}\right)\right] \quad \text{(S5)}$$

where z is the perpendicular distance from a magnet's pole face, $B_r$ denotes the remanence magnetic flux density of 1.186 T, and l, w, h, are the length, width, and height, with 32 mm, 27 mm, and 6 mm, respectively.

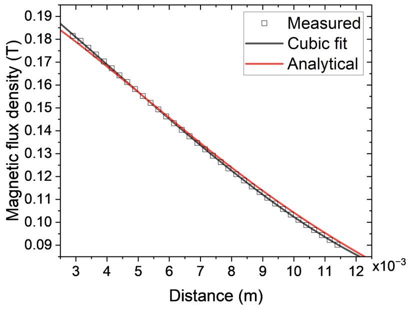


**Figure S3.** Measurement of the magnetic flux density in the sensor plane with respect to the distance of the cubic permanent magnet.

### 6. Sensor chip layout

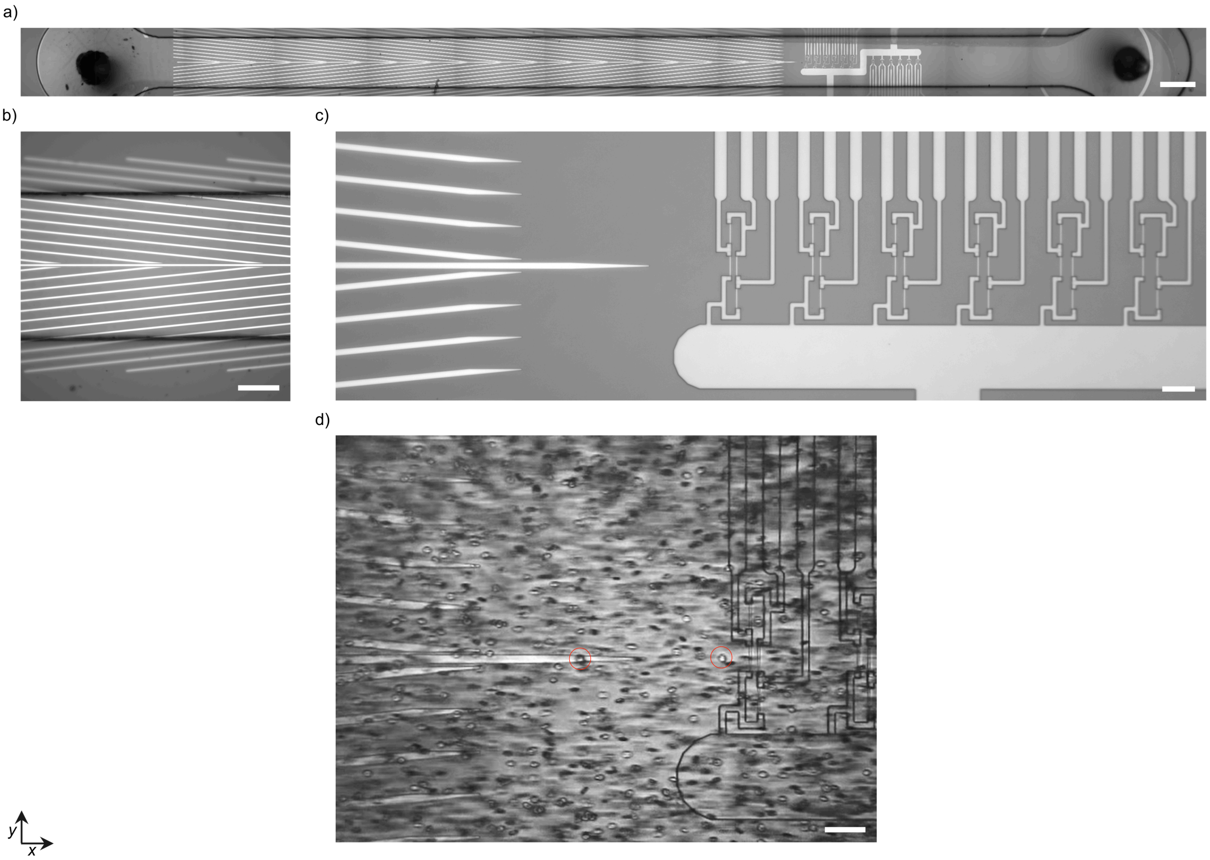


**Figure S4.** Top view of the sensor chip with microfluidic channel. The sample flow is in the *x*-direction. **a)** Panorama image of the sensor chip, including the magnetic rails and GMR sensor arrays assembled with the fluidic channel. The scale bar is 500 µm. **b)** Magnetic rails focus magnetically labeled cells into the channel's center. The scale bar is 200 µm. **c)** The central magnetic rail aims at the GMR sensor array that is configured in Wheatstone half-bridges. The GMR sensor bridge distance within a Wheatstone half-bridge varies between 10 µm and 22 µm, although only bridges with distances between 14 µm and 18 µm were used (3rd to 5th bridge from the left). The scale bar is 50 µm. **d)** The whole blood sample is transported over the sensor. Immunomagnetically labeled $CD14^{+}$ monocytes (circled in red) are focused by the magnetic rails and pass over the GMR sensors, inducing the distinct signal. The scale bar is 50 µm.

### References

Gou, X.-F., Yang, Y., Zheng, X..J., 2004. Applied Mathematics and Mechanics 25, 297-306.
Reisbeck, M., Helou, M.J., Richter, L., Kappes, B., Friedrich, O., Hayden, O., 2016. Sci. Rep. 6, 32838.
